# Comprehensive Transcriptome Annotation of Thousands of HIV-1 Genomes

**DOI:** 10.1101/2025.09.24.675449

**Authors:** Ales Varabyou, Mykhaylo Artamonov, Sophia Cheng, Diane L. Bolton, Steven L. Salzberg, Mihaela Pertea

## Abstract

Alternative splicing in HIV-1 has been a central focus of decades of research, uncovering key mechanisms of viral gene regulation, immune evasion, and therapeutic response – yet, no reference resource has existed to support transcriptome-wide analysis, limiting adoption of modern computational methods. We present HIV Atlas (https://ccb.jhu.edu/HIV_Atlas), the first reference-quality annotation of HIV-1 and SIV transcriptional diversity. We manually curated transcriptomes for HIV-1_HXB2_ and SIV_mac239_ and developed Vira, an automated annotation-transfer method specifically designed to address unique challenges of viral genome biology, to generate high-quality annotations for 2,077 complete HIV-1 genomes. Using the resources presented in our work, we evaluated conservation of splice sites, revealing near-perfect preservation of major donors and acceptors. Furthermore, using several public datasets, we demonstrate how HIV Atlas enhances methodology, improves the quality and novelty of results, and opens novel avenues for research, supporting more accurate and comprehensive analyses of bulk, single-cell, and spatial RNA-seq in HIV-1 studies.

## 1. Main

Human immunodeficiency virus type 1 (HIV-1) remains a major global health burden, with over 40 million people living with HIV and more than 600,000 deaths annually due to HIV-related illnesses^1^. Yet, despite decades of research and effective antiretroviral therapies, viral latency and persistence present ongoing challenges to eradication^2, 3^. In particular, splicing dynamics of HIV-1 play a crucial role in viral replication, pathogenesis, and interaction with the host, and present promising therapeutic targets^4-8^. Due to the diversity of HIV-1 genomes and computational challenges, no comprehensive and consolidated resource exists to systematically catalog this body of knowledge using modern genomic technologies to study transcriptional dynamics of the virus. In this work, we address this gap by presenting the first consolidated atlas of transcriptional diversity for HIV-1 and SIV genomes, alongside an automated framework for extending this resource to new isolates.

HIV-1 is a complex retrovirus that produces its proteins through an elaborate splicing process, enabling the synthesis of 15 protein products from 9 open reading frames (ORFs) in a ∼10 kilobase (kb) RNA genome^9, 10^. After integration of the proviral DNA into the host genome, a single RNA transcript is generated that undergoes alternative splicing, giving rise to three main types of mRNAs: unspliced genomic mRNA that encodes structural and enzymatic proteins (Gag and Pol polyproteins), partially spliced transcripts that produce envelope and accessory proteins (Env, Vif, Vpr, and Vpu), and fully spliced transcripts that produce regulatory proteins (Rev, Tat, and Nef)^11-14^. These splicing events regulate the expression of viral proteins essential for replication, immune evasion, latency, and viral reactivation. RNA sequencing (RNA-seq) has been instrumental in uncovering the complexity of these splicing patterns, illuminating how different transcript isoforms might contribute to viral persistence, pathogenicity, host immune response, and response to treatment^11, 12, 15, 16^. Recent advances in high-throughput sequencing, particularly long-read and single-cell technologies, have made it feasible to capture full-length viral transcripts at unprecedented resolution, further emphasizing the utility of transcriptomic analyses to the study of HIV-1^4, 12, 17^.

Since the inception of RNA sequencing, numerous bioinformatics tools have been developed for efficient and accurate transcriptomic analysis^18^. Novel transcripts are routinely identified in different organisms by assembling short-read and long-read data using tools such as StringTie, IsoQuant, and Bambu^19-22^. However, these computational approaches greatly benefit from, and often require, detailed prior information about trusted gene and transcript models for correct read-to-transcript assignment and quantification^23-25^. In contrast to most model organisms, HIV-1 and SIV lack such foundational resources with no comprehensive or unified transcriptome annotation available for these viruses. This absence limits the application of established RNA-seq protocols to viral datasets, reducing accuracy and interpretability, and introducing inconsistencies across studies. Even when viral RNA is sequenced, results often remain underutilized due to the lack of context provided by a shared reference framework.

Besides opening doors to entirely new analytical frameworks in comparative and functional genomics, the proper incorporation of reference splice junctions into the alignment, assembly and quantification of RNA-seq data has been shown to dramatically improve accuracy, with benefits proportional to the completeness and comprehensiveness of the annotations^22, 26, 27^. In recent years, the rise of single-cell RNA-seq has motivated the adaptation of quantification methods such as Salmon, Kallisto, and STAR to quantify spliced versus unspliced RNA, a task that requires prior knowledge of the transcriptional structure of the genome^28-31^.

This challenge is further exacerbated in HIV-1 and SIV by their extreme sequence diversity, often driven by extensive mutagenesis, hypermutation, recombination, and rapid replication cycles. Nucleotide divergence within a single HIV-1 subtype can range from 15% to 20%, while differences across subtypes can exceed 25%–35%^32, 33^. These mutations affect not only the proteins produced by the provirus but also affect splicing dynamics, potentially altering the transcriptional properties among isolates^34-36^. Such variability complicates the use of a single reference genome and annotation for all studies, highlighting the importance of isolate-specific information. Indeed, it is this very diversity that both necessitates a comprehensive annotation effort and makes it technically challenging. The lack of a standardized transcript catalog is not simply an oversight: it reflects the profound biological complexity of these viruses and the limits of existing annotation frameworks.

While multiple reports of HIV-1 and SIV splicing exist, this information is rarely accessible in a standardized format suitable for computational analysis, limiting the analysis of RNA sequencing datasets from human and animal infection models. Some studies provide viral transcriptome annotations along with corresponding GTF files^11, 37, 38^, but these are often non-standardized, lack detail, and rarely extend beyond specific isolates used in individual studies. Furthermore, constructing such resources requires substantial manual effort, specialized expertise, and is prone to errors. Other studies make use of known splicing information internally to improve HIV-specific computational workflows^39^, yet the included information is limited to the methods used and inaccessible for broader computational use. Overall, the current lack of a gold standard resource complicates and diminishes the quality of downstream comparative analyses across study.

In this work, we address the critical gaps outlined above by presenting HIV Atlas, the first comprehensive and consolidated dataset of transcriptional diversity for HIV-1 and SIV genomes, along with an automated framework for extending this resource to new isolates. Our atlas captures transcriptional diversity across thousands of publicly available viral isolates, spanning a broad range of geographic origins and all major and minor subtypes, which we systematically annotated using Vira, a novel computational method specifically designed to transfer transcript annotations across diverse viral genomes. By leveraging extensive literature, integrating novel computational strategies, and validating against public datasets, we provide a robust resource that encapsulates the dynamic splicing architecture of these viruses. Our resource enables numerous applications, including the expansion of targets for sensitive probe designs, the development of better predictive models informed by precise splicing information, new studies on RNA secondary structure and translation regulation, a deeper understanding of host-pathogen interactions, and the improvement of phylogenetic models.

## 2. Results

### 2.1. Manual Construction of HIV-1_HXB2_ Reference Genome Annotation

We chose HIV-1_HXB2_ (GenBank accession K03455.1)^40^ as our reference genome due to its frequent use as the reference strain and coordinate standard in functional and comparative HIV studies, and because it already has a complete annotation of open reading frames (ORFs)^41, 42^. We constructed the base annotation of the HIV-1_HXB2_ genome following established conventions^13, 43^.There has been a recent push for human transcript catalogs to be more rigorous, i.e., to exclude transcripts with weaker evidence of function^27, 44, 45^, because overly inclusive annotation can lead to a bias towards noisy or invalid transcripts. In contrast, a conservative annotation database allows for novel isoforms to be discovered and evaluated carefully. Thus, for the base annotation of HIV-1, we chose to include all previously described conserved splice sites^13^ and all transcripts that contain only these sites.

To annotate the HIV-1_HXB2_ isolate, we incorporated information from prior experimental studies. In particular, we utilized the transcript information from the Ocwieja report, which focused on the analysis of the HIV-1_89.6_ isolate (GenBank accession U39362.2)^46^. To transfer these annotations to the HIV-1_HXB2_ genome sequence, we constructed a map of the coordinates of the major and minor splice donor and acceptor sites between the two sequences. Using the flanking exonic and intronic sequence for each of the donor and acceptor sites on the HIV-1_89.6_ genome as reported by Ocwieja et al., we manually identified the corresponding sites on the HIV-1_HXB2_ genome (Table 1). We then proceeded to reconstruct all previously-described transcript sequences using the donor and acceptor sites as reported by the authors. Only transcripts compatible with their respective proteins were included in the final catalog (Supp. Table 1). During the reconstruction of each transcript on the HIV-1_HXB2_ genome, the sequence of exon coordinates was verified for compatibility with the corresponding annotated protein of that transcript in GenBank (Supp. Table 2). During this process, we found that three transcripts from the Ocwieja et al. report were incompatible with the corresponding protein because they lacked the required D4-A7 splice junction.

**Table 1.**
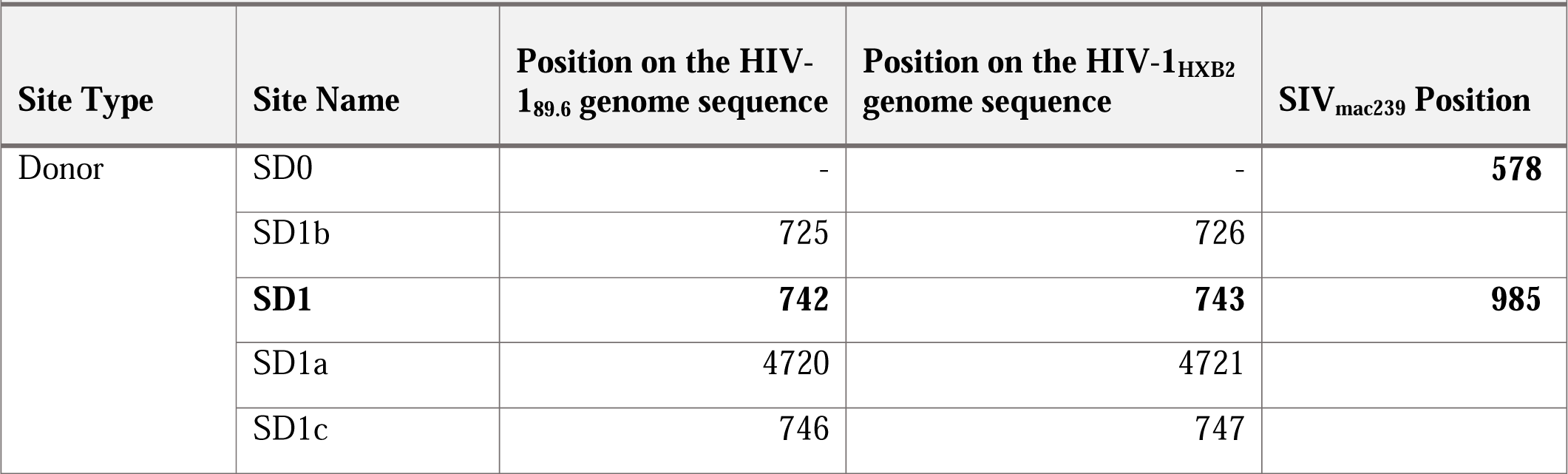
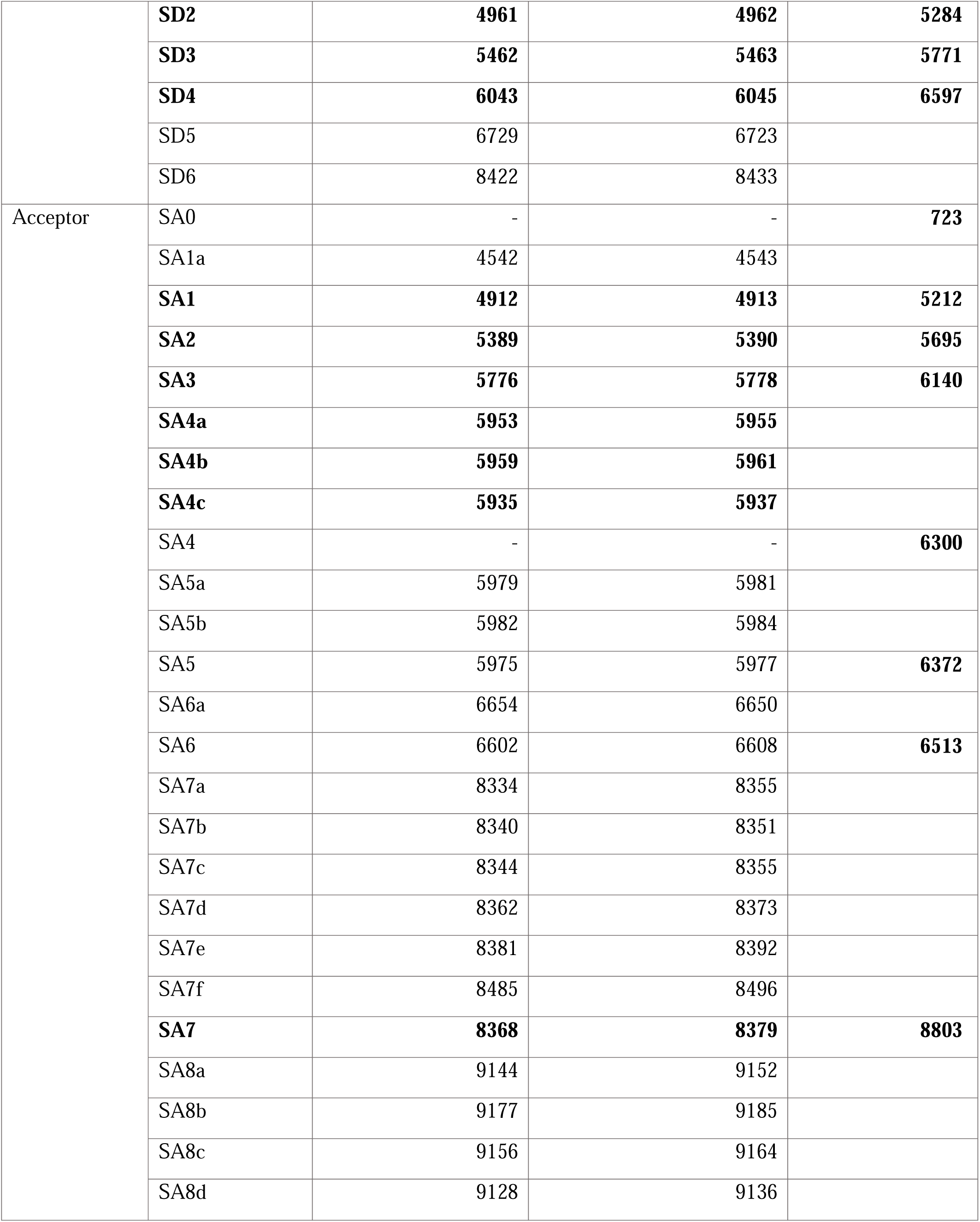
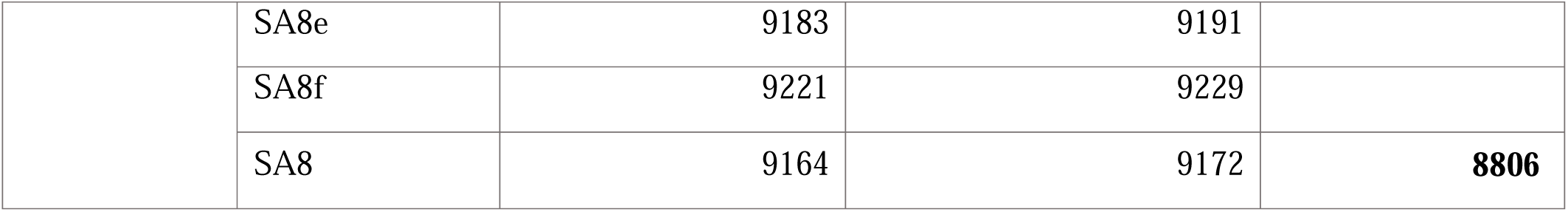
Splice donor and acceptor site positions in HIV-1 and SIV annotations, following the nomenclature from Ocwieja et al. for the HIV-1 annotation and Emery et al.^48^ for the SIV_mac239_ annotation. Donor positions represent the last exonic nucleotide before the splice site, while acceptor positions represent the first exonic nucleotide after the splice site. The SIV_mac239_ genome sequence used is the LANL variant with the 256 bp 5’ host integration flanking region removed. Positions in bold are included in the reference catalog.

| Site Type | Site Name | Position on the HIV-1<sub>89.6</sub> genome sequence | Position on the HIV-1<sub>HXB2</sub> genome sequence | SIV<sub>mac239</sub> Position |
| --- | --- | --- | --- | --- |
| Donor | SD0 | - | - | **578** |
|  | SD1b | 725 | 726 |  |
|  | **SD1** | **742** | **743** | **985** |
|  | SD1a | 4720 | 4721 |  |
|  | SD1c | 746 | 747 |  |
|  | **SD2** | **4961** | **4962** | **5284** |
|  | **SD3** | **5462** | **5463** | **5771** |
|  | **SD4** | **6043** | **6045** | **6597** |
|  | SD5 | 6729 | 6723 |  |
|  | SD6 | 8422 | 8433 |  |
| Acceptor | SA0 | - | - | **723** |
|  | SA1a | 4542 | 4543 |  |
|  | **SA1** | **4912** | **4913** | **5212** |
|  | **SA2** | **5389** | **5390** | **5695** |
|  | **SA3** | **5776** | **5778** | **6140** |
|  | **SA4a** | **5953** | **5955** |  |
|  | **SA4b** | **5959** | **5961** |  |
|  | **SA4c** | **5935** | **5937** |  |
|  | SA4 | - | - | **6300** |
|  | SA5a | 5979 | 5981 |  |
|  | SA5b | 5982 | 5984 |  |
|  | SA5 | 5975 | 5977 | **6372** |
|  | SA6a | 6654 | 6650 |  |
|  | SA6 | 6602 | 6608 | **6513** |
|  | SA7a | 8334 | 8355 |  |
|  | SA7b | 8340 | 8351 |  |
|  | SA7c | 8344 | 8355 |  |
|  | SA7d | 8362 | 8373 |  |
|  | SA7e | 8381 | 8392 |  |
|  | SA7f | 8485 | 8496 |  |
|  | **SA7** | **8368** | **8379** | **8803** |
|  | SA8a | 9144 | 9152 |  |
|  | SA8b | 9177 | 9185 |  |
|  | SA8c | 9156 | 9164 |  |
|  | SA8d | 9128 | 9136 |  |
|  | SA8e | 9183 | 9191 |  |
|  | SA8f | 9221 | 9229 |  |
|  | SA8 | 9164 | 9172 | **8806** |

In HIV-1 and SIV genomes, Env and Vpu proteins are translated from multiple bicistronic mRNAs^47^. However, representing polycistronic transcripts is challenging in GTF and GFF formats because those formats were designed for monocistronic gene models, and require each coding transcript to have a single protein assigned. Deviations from this format can cause malfunctions in downstream software utilities. Because polycistronic cases are very rare in plant and animal genomes, annotators of those species have adopted a practice of duplicating the transcript for each protein in a polycistronic locus. However, that solution inherently interferes with transcript-level expression analysis, because RNA-seq reads will map ambiguously to the two identical copies. To avoid sequence duplication and issues during quantification, we opted to represent all bicistronic transcripts under the same gene identifier, simplifying the analysis. To preserve protein information in the annotation, we annotated the Vpu protein for one of the bicistronic mRNAs at the locus and Env protein for all other transcripts.

On the other hand, the Gag/Pol polyprotein is translated from an unspliced mRNA, which prevents us from using the same strategy as we did for the Env and Vpu bicistronic mRNAs. Instead, we had to use the same transcript annotation with different coding regions annotated for the two proteins, thus resulting in a duplication of the transcript sequence.

As a reference resource, our annotation does not attempt to account for every known exception or molecular mechanism involved in HIV-1 transcriptional and translational regulation. While, numerous exceptions have been documented, including cryptic donor and acceptor sites, isolate-specific splicing events, condition-specific transcripts, and very short ORFs^5, 13, 48-50^, our primary goal is to provide a consistent and reliable control set for a wide range of future studies.

### 2.2. Reference Annotation for the SIV Genome

The SIV atlas was constructed similarly to HIV-1, but because this virus is much less well-studied, we utilized additional resources from short-read sequencing data to create the database. The full annotation was completed as part of an upcoming SIV integration site study.

### 2.3. Data Collection

As of 2022, 5,381 complete HIV-1 sequences were available in the latest LANL HIV Sequence Compendium, an annual publication from the Los Alamos National Laboratory (LANL) that compiles and analyzes HIV genetic sequences collected from around the world. This curated set of high-quality complete genome assemblies is commonly used to compare newly sequenced viruses with existing strains and to study viral evolution over time. The collection itself represents a valuable aggregation of viral diversity and is often used to study the variation of the HIV genome, and to track changes in the HIV genome that affect vaccine targets or antiretroviral drug resistance^51^. Additionally, many widely utilized HIV-1 isolates are included in this dataset. To facilitate downstream analyses and enable more comprehensive studies of HIV genomic variation, we supplemented the genome assemblies and protein annotations provided in the compendium and corresponding GenBank annotations with complete transcript structures for each genome, thereby enhancing the utility of HIV Atlas as a reference resource for a broad range of applications.

To this end, we obtained the 2022 revision of the LANL HIV Compendium^52^, including 5,359 complete sequences, omitting 22 accessions for which we encountered errors when fetching the data. For each of the accessions included in the compendium, we obtained the corresponding sequence and GTF annotation through GenBank ^53^ by utilizing a custom web crawler.

We applied specific inclusion criteria to determine which genome sequences to retain in our compendium set. The most critical requirement was sequence completeness. While the LANL compendium defines completeness based on the presence of all protein-coding regions, this standard does not always guarantee inclusion of all non-coding regions which are essential for our computational framework. In particular, because splice donor 1 (SD1) is located upstream of the Gag region, we refined the definition of completeness to require the presence of this site. To test for this added requirement, we used minimap2^54^ with the ONT preset to align the HIV-1_HXB2_ (GenBank accession K03455.1) genome sequence, which we used as our reference, to each of the downloaded compendium sequences. We then confirmed that all reference splice donor and acceptor sites, as well as CDS regions, were covered in the alignments. This filtering step eliminated 2,289 accessions from the initial set, leaving 3,070 genomes for further analysis. While our approach could also be applied to incomplete genomes, we limited this work to complete genomes to ensure HIV Atlas serves as a comprehensive and reliable reference resource.

Lastly, all GenBank protein annotations initially fetched in the GFF3 format were standardized using gffread^55^ to make sure that the formatting was uniform across all genomes. We also obtained metadata for all complete genomes from the LANL database and appended key fields, namely subtypes, years, geographic region and sequence name, to our existing metadata, which can be used to search and navigate the atlas’s web interface.

### 2.4. Automated HIV/SIV genome annotation with Vira

To facilitate the transfer of our manually curated reference annotations of the HIV-1 (HIV-1_HXB2_) and SIV (SIV_mac239_) genomes onto genome assemblies of other isolates, we designed a fully automated computational protocol, Vira (see Methods). While methods have been developed for annotating genomes entirely de novo or by transferring reference annotations from closely related species^56-59^, none were designed for the specific challenges presented by the HIV and SIV genomes, with their high degree of sequence divergence, hypermutation, and recombination.

We first assessed the accuracy of Vira by annotating two viral isolates with previously characterized splicing sites and comparing our automated annotations to those reported in the original studies. Having validated the protocol, we applied our software to perform annotation transfer from the HIV-1_HXB2_ onto thousands of curated HIV-1 genomes. Using these annotations, we then examined the conservation of major splice donor and acceptor sites across diverse HIV-1 isolates. Lastly, we demonstrate how the resources from our study can be seamlessly integrated to enhance RNA-seq analyses in studies of both HIV-1 and SIV.

#### 2.4.1. Annotation inconsistencies in HIV-1 genomes

Because the original study by Ocwieja et al. used the HIV-1_89.6_ genome as the reference for their catalog of splicing events, we wanted to test our protocol on that genome first, confirming the validity of our automated approach and the consistency of the resulting annotation with the original study. To our surprise, our software detected an error due to incompatibility with the HIV-1_89.6_ protein annotation in GenBank. To investigate this issue, we compared the annotation provided by GenBank to what our software generated and found that the SD4-SA7 junction was erroneously annotated. As illustrated in Figure 1.A, the GenBank annotation (shown in red) has a 1-nucleotide truncation at the end of the coding exon upstream of the junction at the SD4 donor site, which is compensated by a 2-base truncation of the downstream coding exon at the SA7 site. These truncations, which remove an amino acid from the Tat and Rev proteins, also result in splice sites that fail to respect the canonical GT-AG splicing signals, which our annotation (in orange) uses properly.

**Figure 1.**
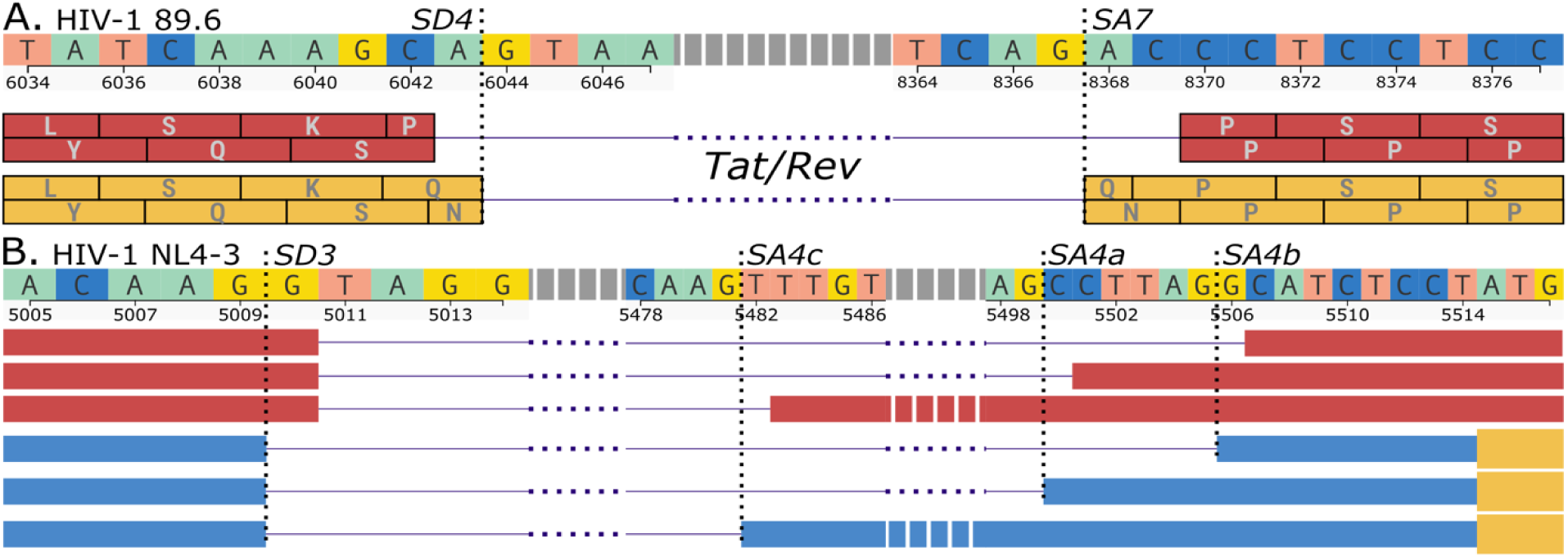
Erroneous HIV-1 annotations in circulating research. A. Inconsistency in SD4 donor and SA7 acceptor splice site annotations in the coding sequence of the GenBank HIV-1_89.6_ (U39362.2) annotation. The GenBank annotation (shown in red) incorrectly positions the SD4 donor site one base upstream of the canonical GT dinucleotide and the SA7 acceptor site two bases downstream of the canonical AG dinucleotide. This erroneous annotation results in the truncation of the encoded protein sequence by one complete amino acid. Our suggested corrected annotation (in orange) is shown below the GenBank annotation. The image shows the NCBI Genome Viewer interface, with intronic sequences represented by dashed elements for clarity. B. Inconsistencies in the non-coding annotation of HIV-1_NL4-3_ molecular clone. An off-by-one error in the genome annotation around the donor and acceptor sites (SD3, SA4a,b,c illustrated here) of the HIV-1_NL4-3_ genome is illustrated with a zoomed-in view of the annotation provided by Bohn et al.^37^. The upper tracks (in red) illustrate a subset of the junctions from the Bohn et al. annotation, with shifted donor and acceptor sites that do not use the canonical GT/AG dinucleotides. The bottom tracks (in blue) show correct placement of the splice junctions and exons, as provided by the Vira package.

Having identified the error, we repeated our automated annotation using the corrected HIV-1_HXB2_ protein set as the reference. The updated results were consistent with the original report from Ocwieja et al. for all donor and acceptor sites, including the SD4–SA7 junction, as well as for the overall protein annotation.

#### 2.4.2. Validation of Vira Annotations against Existing Manual HIV-1 Annotations

To further validate our protocol, we searched for another HIV-1 isolate that had high-quality, manually curated annotation. Such resources are rare, but we identified a single study that provided a GTF annotation of their target HIV isolate^37^. This resource, compiled in part from the same datasets we used, included a detailed manually curated annotation accounting for all major donor and acceptor sites, as well as several cryptic sites. Upon closer examination, we identified several inconsistencies in this annotation, likely stemming from coordinate conversion errors. The first issue was a 1-bp shift in splice junction coordinates upstream of donor sites and another 1-bp shirt downstream of acceptor sites, as illustrated in the SD3 to SA4b,a,c sites in Figure 1.B. Additionally, we found that transcript sequences were truncated at the 3’ end due to trimming of the sequences to the PCR primer ends – an intentional choice by the original authors that nonetheless limits the usability of this auxiliary resource. Lastly, this manually curated resource did not include ORF annotations for any of the transcripts.

We applied our annotation protocol, Vira, to annotate the same genome used in the study. Although direct comparison was challenging due to the coordinate discrepancies in the original annotation, manual validation to account for these errors confirmed that Vira annotations agreed perfectly with all donor and acceptor sites reported in the study. Notably, Vira successfully recovered the unannotated 3’ region missing from the original annotation due to the intentional trimming of the sequences to the PCR primer ends, including correct recovery of the missing D4-A7 junction, with correct placement of donor and acceptor sites at the canonical GT-AG dinucleotide flanks.

It is important to note that the coordinate discrepancies in this use case likely did not affect the original study’s analysis, which primarily relied on long-read data. However, this example illustrates how such issues could be avoided through access to a standardized and reliable annotation resource such as the one we propose.

#### 2.4.3. Annotation of thousands of complete HIV-1 genome assemblies with Vira

After validating the quality of the annotations produced by our method, we used the Vira software to perform annotation transfer from the HIV-1_HXB2_ onto the 3,070 genomes that remained after our initial filtering to remove incomplete genomes from the LANL Compendium. For 2,077 of these genomes, Vira transferred the annotation with no errors. Several issues prevented the system from completing annotation of the remaining genomes, with the majority of problems caused by minimap2 alignments that failed to capture one or more splice sites, most often beginning downstream of the SD1 site. These failures may reflect mutations at canonical splice sites, masked sequence in the regions flanking those sites, or other systematic inconsistencies in the assemblies that prevented accurate transcript alignment. In rarer cases, Vira was unable to complete annotation transfer because of disagreements between splice sites inferred from the reference transcriptome alignments and those reported in available guide annotations. Such discrepancies could stem from alignment errors or inconsistencies in the guide annotations themselves, and would require additional manual curation of the genome sequences alongside closer examination of the protein and partial transcript alignments to resolve. For the 2,077 successfully annotated genomes, we proceeded with downstream comparative analyses.

#### 2.4.4. Conservation of Donor and Acceptor Sites across HIV-1 genomes

Analysis of splice site sequences across the 2,077 HIV-1 genomes annotated by Vira revealed strong conservation of the major donor and acceptor sites (Figure 2). All analyzed donor sites and seven of eight acceptor sites exhibited perfect conservation of the canonical GT and AG dinucleotides, respectively. One acceptor site, at position 5,937, showed slight variation, in which a small fraction of the isolates contained the non-canonical dinucleotide AC rather than AG. Overall, splice sites displayed significantly higher conservation than the rest of the genome (p < 0.001, Mann-Whitney U test). While splice sites showed near-perfect conservation (entropy: μ = 0.0002, σ = 0.0008), the remainder of the genome exhibited substantially higher sequence variation (entropy: μ = 0.3889, σ = 0.3686).

**Figure 2.**
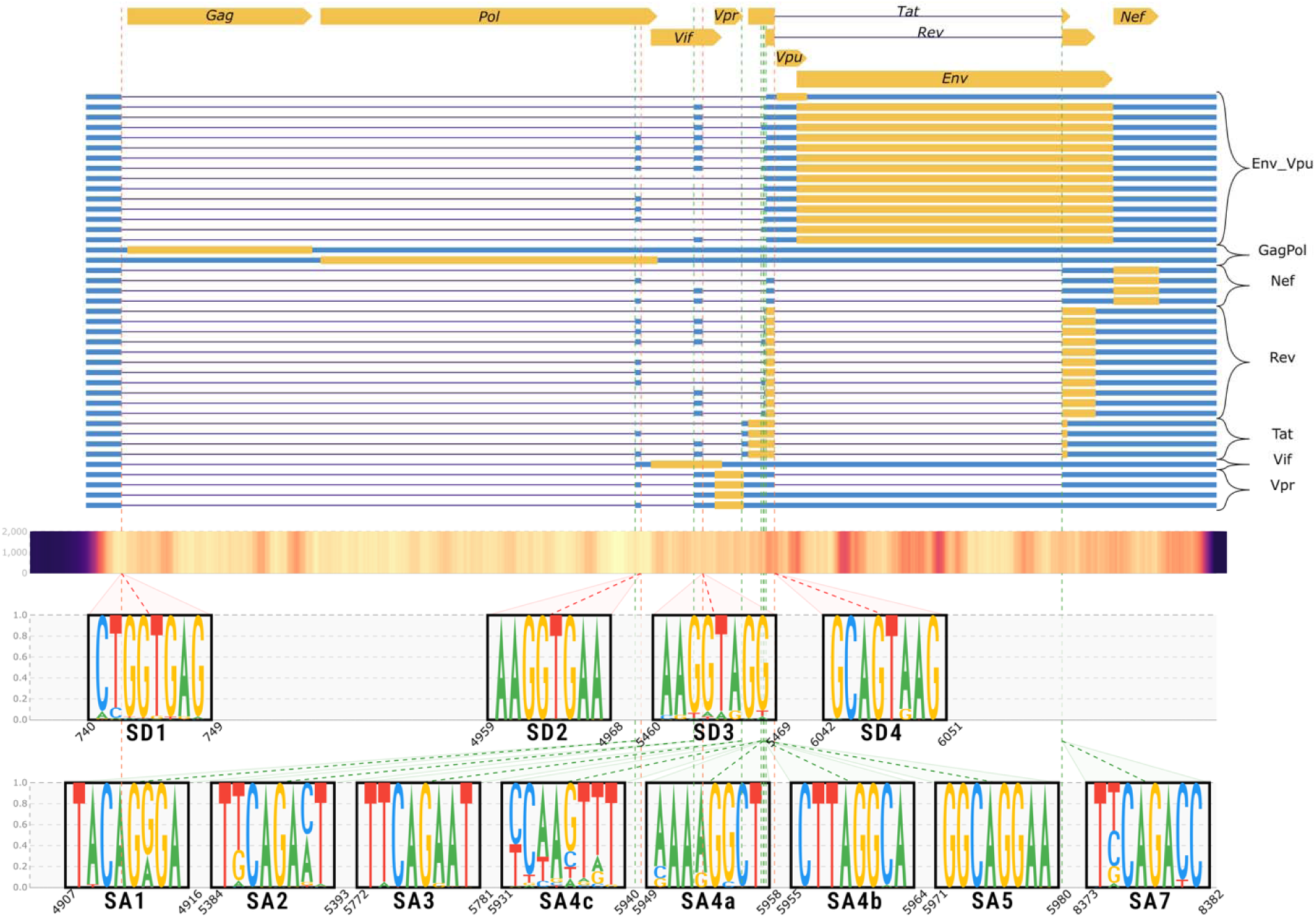
Atlas of HIV alternative splicing and conservation of annotated splice donor and acceptor sites, mapped onto the HIV-1_HXB2_ (K03455.1) genome coordinates. The top panel depicts the nine reference HIV ORFs. Below, a diagram illustrates the 41 alternative splice forms annotated for the HIV-1_HXB2_ isolate, grouped by gene. Each alternative splicing event includes non-coding transcribed sequences (blue), proteins (yellow), and introns (black lines). Middle panel: a heatmap of conservation scores across the HIV-1_HXB2_ genome, derived from the Vira alignment of 2,077 annotated genomes, with darker shades indicating lower conservation. Dashed lines mark donor (red) and acceptor (green) sites across all panels. Bottom: close-up views of each annotated splice site display sequence logos, including the consensus dinucleotide (AG/GT) and three flanking bases on each side.

### 2.5. HIV and SIV Annotation Web Resource

To present our results to the broader scientific community and facilitate easy exploration and usage of our dataset, we constructed an interactive web resource called HIV Atlas. HIV Atlas provides a user-friendly interface for searching over the database of HIV-1 and SIV genomes and their annotations, which are hosted on GitHub for transparent and robust version control. For each genome, the database provides standardized and searchable metadata consisting of: (1) country of origin, (2) year of collection, (3) subtype, (4) description from the original submission, (5) annotation score, and (6) GenBank accession identifier. The web application integrates the JBrowse 2 genome browser^60^ with a streamlined view of the genome sequence and annotation, providing quick and easy way to explore and compare genomes (Figure 3).

**Figure 3.**
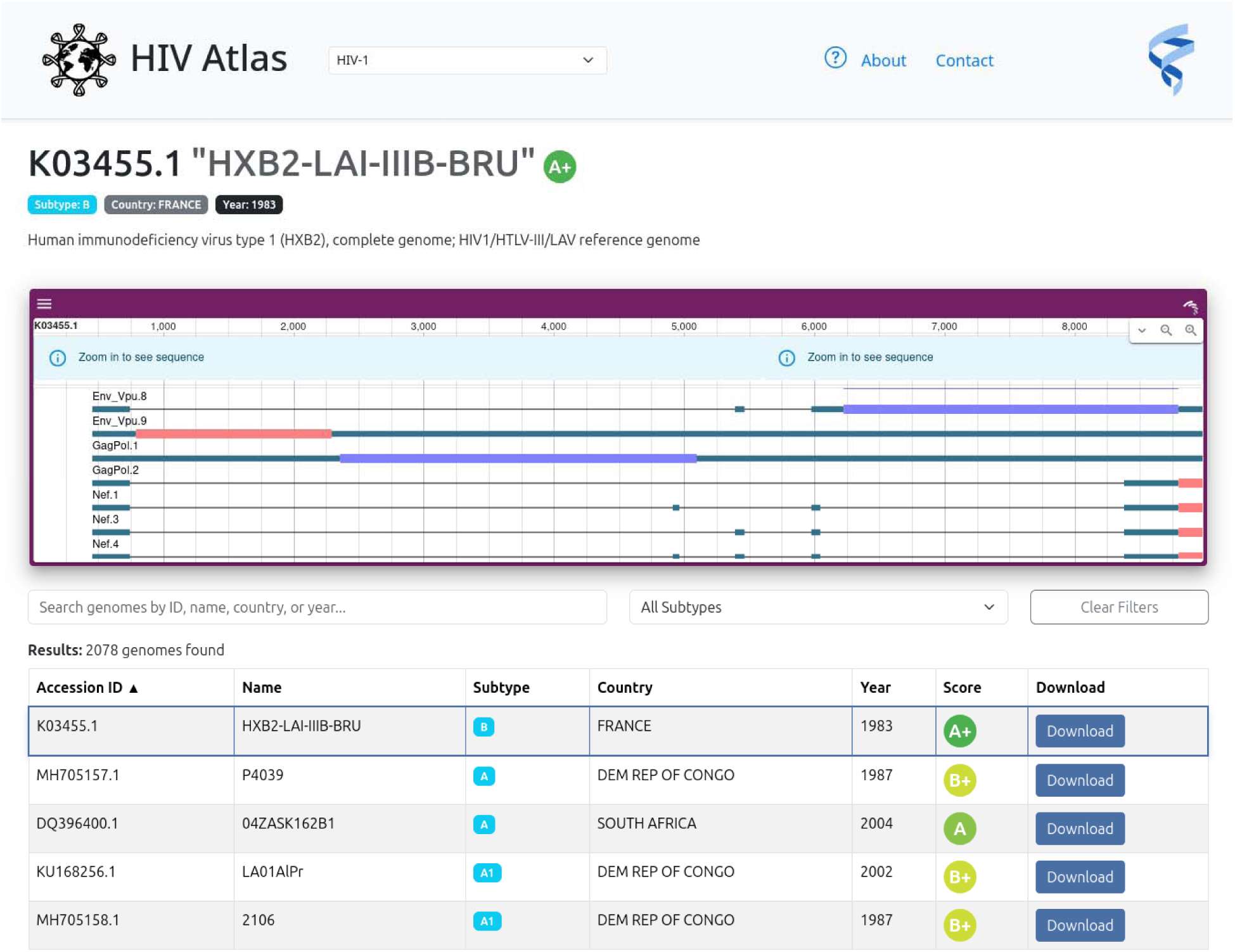
Screenshot of the HIV Atlas web interface. The web application integrates a filter and search interface with matching results displayed in a table along with minimal metadata, such as the common name, subtype, country and year of collection and annotation score. Each of the genomes in the atlas can be explored directly on the web page via the JBrowse2 genome browser together with a full set of transcripts and protein available.

### 2.6. Enhanced Analysis of Public HIV and SIV RNA-seq Data

To demonstrate the utility of our atlas and annotation protocols in analyzing HIV and SIV datasets, we conducted several proof-of-concept experiments focused on quantifying transcriptional differences using public datasets. While not exhaustive, these examples highlight the most fundamental and broadly applicable features of our resource: transcript assembly, classification, and quantification for both established and novel viral isolates, using both short and long-read sequencing data with varying levels of noise and errors.

First, we analyzed SmartSeq single-cell RNA-seq data from SIV-infected rhesus macaque CD4^+^ T cells^17^, showcasing how annotations obtained directly from the HIV Atlas improve detection of differences in transcriptional dynamics between experimental conditions. Second, we examined long-read sequencing data generated with the Oxford Nanopore Technology (ONT) from HIV-1_NL4-3_ infected human CD4^+^ T cells^12^, illustrating how our software can be utilized to annotate novel HIV-1 genome assemblies and aid in accurate transcript reconstruction, discovery and quantification. Together, these examples highlight the versatility and practical value of our annotation resources across diverse experimental contexts.

#### 2.6.1. Transcriptional differences in SIV expression during acute infection

First, we illustrate the use of HIV Atlas on full-length, single-cell SmartSeq RNA-seq data from SIV^17^. Because our goal was to illustrate the application of the SIV annotation to the analysis of the viral transcriptome, we focused exclusively on viral expression and ignored the host gene expression. For our investigation, we selected a subset of 149 samples comprised of infected CD4^+^ T cells actively transcribing SIV, defined in the original study by the presence of spliced viral reads (svRNA). The high svRNA content of the samples, averaging 7-10% of the total reads, enabled detailed characterization of transcriptional diversity. The dataset also included measurements from two infected host macaques: memory CD4^+^ T cells (CD95^+^) from a mesenteric lymph node (SIV_mac251_-infected animal AY69, n=89 cells) and from peripheral blood (SIV_mac239X_-infected animal T034, n=60 cells) providing us with two distinct groups of samples to analyze and compare. In this demonstration, we were particularly interested in leveraging the rich RNA-seq data to assess the extent to which host- or tissue-specific factors may influence viral splicing. By supplementing the analysis with the transcriptome annotations we set out to compare transcriptional profiles between the two groups in terms of differential transcript expression, as well as to exploit the full-length nature of the dataset to search for previously uncharacterized isoforms.

To begin our analysis, we first used the HIV Atlas web interface to locate the genome and annotation corresponding to the SIV_mac239_ isolate from the original study. To avoid erroneous mapping of host reads present in the data to the viral genome, we prepared a joint reference by combining the macaque MMul10 genome and its corresponding Ensembl annotation^61^ with the SIV_mac239_ reference genome and annotation. For alignment of the sequencing data, we chose HISAT2 ^62, 63^, although other aligners such as STAR^64^ or minimap2^54^ could be used instead. The HISAT2 index was constructed using the combined reference genome and augmented with the splice junction and exon information extracted from the joint annotation. This approach improved alignment accuracy by reducing multimapping and enhancing splice junction detection, particularly near short exons^62, 65^. For each animal, we aggregated the aligned samples using TieBrush and extracted the combined coverage and junction data using TieCov^66^ to use in downstream comparisons (Figure 4). Read counts for junctions present in the reference annotation were obtained for each animal from the TieCov aggregations.

**Figure 4.**
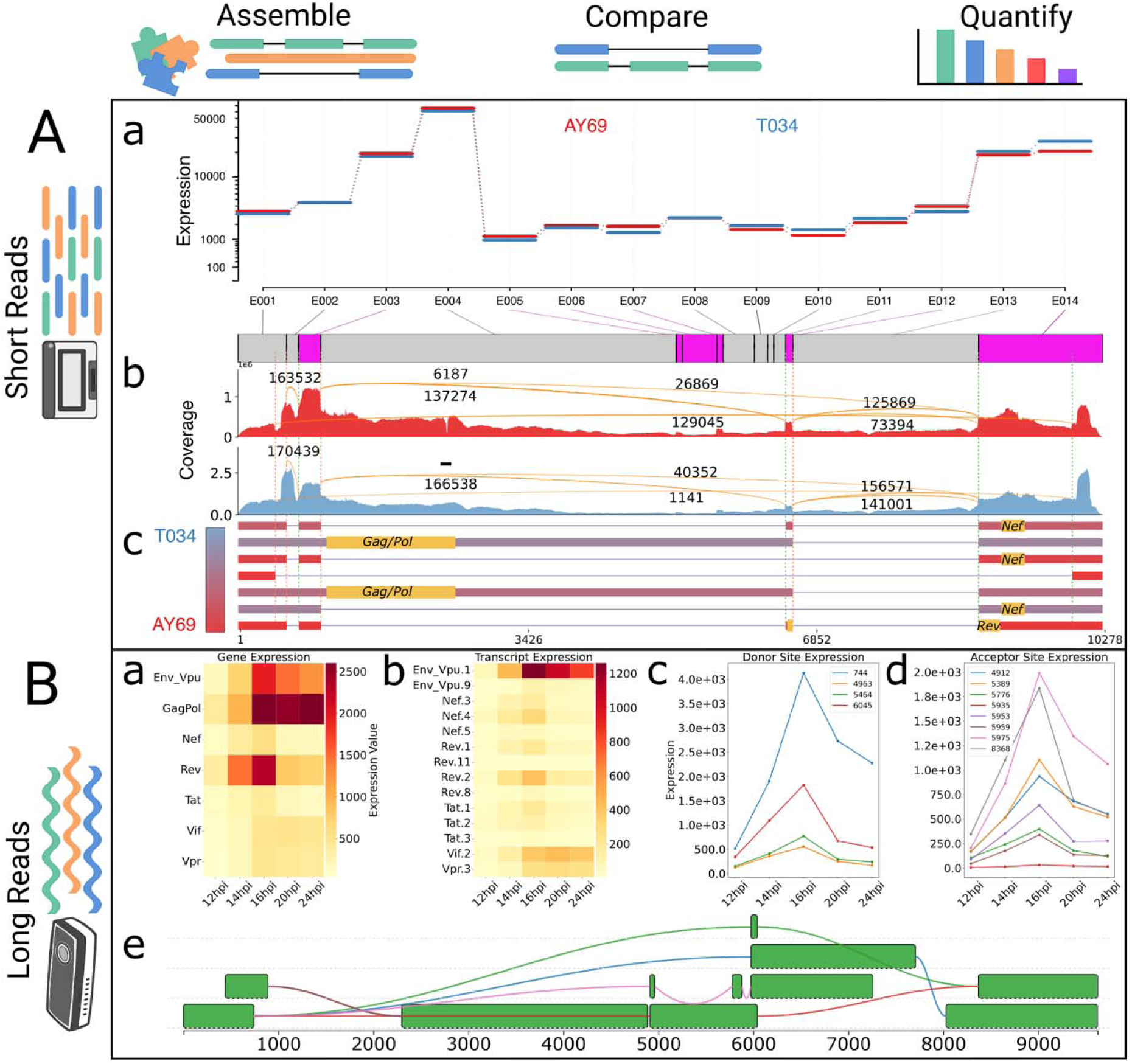
Schematic illustrating experimental applications of the HIV Atlas. Panel (A): Results of analysis of the Jiménez et al dataset^17^ of single-cell SIV expression generated using short-read Illumina sequencing. a. DEXSeq SIV exon expression comparison between sample groups from two macaques. Exons tested for differential usage are numbered E001 through and E014. Exons found to have statistically significant differential usage between the two groups are shown in pink and those without significant variation in grey. b. Coverage profiles of the SIV_mac239_ expression for the two animals. Included junctions and corresponding junction read counts correspond to the junctions observed in the highly expressed novel transcripts shown in sub-panel c below and are aligned with the DEXSeq exons in panel a above. c. Novel transcripts detected in >=10% of the samples, colored on a gradient according to the fraction of samples from each group. Panel B: Results of the tests performed using the long-read Oxford Nanopore data from Quang et al^12^. a. and b. Heatmaps showing changes in gene and transcript expression over hours post infection (hpi), based on read counts. c. and d. Long-reads supporting each donor and acceptor sites. e. Splice graph of the novel isoforms detected by IsoQuant^21^ from the data.

Multiple methods exist for classifying, quantifying, and discovering alternative transcription events from short-read RNA-seq datasets. In this example, we opted to follow the widely adopted StringTie protocol to classify reference transcripts and assemble novel transcripts in the data^22^. For each sample, we assembled and quantified the transcriptome using StringTie2. For each animal, sample-level transcriptome assemblies were merged using gffcompare^55^ to obtain two complete transcriptomes and separate quantification measures. To identify novel alternative splicing events across the entire dataset, we then merged the assemblies from both animals into a single comprehensive transcriptome. All novel transcripts were subsequently annotated with proteins using ORFanage^67^, which was used to assign the most upstream complete reference protein identifiable for each assembled transcript.

As shown in Figure 4.A.b, both sample groups, each corresponding to one of the two animals, had full genome coverage, with pronounced peaks corresponding to annotated exons. Several differences between the two animals were observed based on our initial analysis, namely a slight increase in the relative abundance of the first translated Rev protein exon at coordinates 6,513 – 6,597 (DEXSeq exon E012) in AY69 sample group compared to T034. Additional differences were observed near the SD0 splice donor site, with a marked shift in dominant peaks in the 5’ LTR region (E001 – E003), between the two groups.

To investigate these differences further, we analyzed the aligned samples using DEXSeq for differential exon usage^68^. While we chose DEXSeq for this demonstration, other statistical frameworks for differential gene expression (DGE), transcript expression (DTE), and transcript usage (DTU) analysis^69-75^ could also be utilized for a more in-depth scrutiny of the data, using our provided annotation in a similar manner as we did with DEXSeq. As shown in the DEXSeq results (Figure 4.A.a), the analysis confirmed our initial observations, identifying five exons with significantly elevated usage. Additionally, a switch in exon usage was detected at the junction between exons E007 and E008, likely indicating slightly elevated levels of multiply spliced svRNA in the T034 sample group.

While the DEXSeq analysis identified significant shifts in exon usage between the two groups, it did not account for the clear differences in expression and splicing observed in the TieBrush alignment data near the SD0-SA0 junction at the 5’ LTR. To investigate these differences further, we examined the most abundant novel transcripts for each sample group. All transcripts were first classified against the reference annotation with gffcompare, and novel transcripts were identified as those lacking a complete intron-chain match with any reference transcript. These novel transcripts were assigned to genes based on the ORFs annotated by ORFanage and were sorted by the number of supporting samples and their expression levels. We found 7 novel transcripts that were present in more than 10% of the samples. Notably, we observed one assembled transcript connecting the 5’ and 3’ LTR regions with a novel splice junction that was expressed exclusively in the AY69 group of samples, which explained the coverage and splicing differences we observed from the TieBrush-ed alignment data.

#### 2.6.2. Annotation of HIV-1_NL4-3_ Molecular Clone and Analysis of ONT long-read cDNA dataset

For the second application, we focused on analyzing long-read sequencing data, which has become increasingly popular for studying transcriptional activity in HIV-1^12, 76^. Oxford Nanopore Technology (ONT) sequencing machines can easily generate reads spanning the entire length of viral mRNAs, offering a comprehensive view of transcript structures. However, high sequencing error rates, further exacerbated by the hypermutability of the virus, make accurate analysis challenging, especially around the splice junctions. Alignment tools optimized for longer reads, such as minimap2, can effectively utilize prior splicing junction annotations to guide and refine challenging alignments.

For this analysis, we selected a particularly suitable dataset generated by Quang et al^12^ as part of their efforts to study longitudinal transcriptional changes in HIV-1 using ONT sequencing. We focused on a subset of the data that tracks changes in svRNA expression in donor CD4^+^ T cells infected in vitro with HIV-1_NL4-3_ and sequenced at intervals of 12, 14, 16, 20 and 24 hours post infection (hpi).

As in our previous analysis, we began by creating a joint index, this time by combining the human and HIV-1_NL4-3_ genomes. However, to our surprise, the commonly used HIV-1_NL4-3_ genome was not included in the list of accessions we obtained from the HIV-1 LANL 2022 Compendium. This presented us with an opportunity to test how the Vira package can handle the annotation of a novel isolate to prepare it for downstream analysis. We used default parameters for Vira to annotate the HIV-1_NL4-3_ reference genome guided by the HIV-1_HXB2_ reference protein annotation from GenBank, fully replicating the protocol we employed for the creation of the HIV Atlas. The resulting HIV-1_NL4-3_ reference annotation was then combined with the CHESS annotation^27^ for the GRCh38 human genome^77^ to complete the creation of a reference dataset for this experiment. The reference dataset was used to create a minimap2 index augmented with the junction data.

We downloaded 5 samples corresponding to the 5 timepoints post-infection and aligned them to the joint reference genome index using the recommended ONT preset for cDNA reads, following the protocol described by Quang et al. While StringTie2 can be effectively applied to both short and long read data, here we chose to use IsoQuant^21^, a method that has recently gained popularity in HIV-1 transcriptional studies^11, 37^. IsoQuant is specifically designed to reconstruct and quantify transcript models from long-read RNA-seq data and incorporates correction techniques tailored to the noisy ONT data. We ran IsoQuant with default parameters, obtaining both assembled novel transcript models as well as quantifications of the reference transcriptome.

Our observations of transcriptional dynamics during the 12-hour post-infection time course closely mirrored the results reported in the original publication, showing peaks at the 14–16-hour mark with a gradual decline thereafter (Figure 4.B.a-b). To further replicate the original findings, we aggregated read counts based on transcript quantifications to derive combined counts for each donor and acceptor site (Figure 4.B.c-d). As expected, these results closely resembled the patterns observed in the original study and reflected the same trajectories of mRNA abundance. Interestingly, our analysis of the novel transcript models reported by IsoQuant, revealed several unannotated splice junctions and exons absent from the original study’s findings (Figure 4.B.e).

## 3. Conclusion

Despite decades of progress in HIV research, the lack of a comprehensive and standardized transcriptome annotation for HIV-1 and SIV remains an open problem hindering adoption of many state-of-the-art RNA-seq technologies in the field. The unusual diversity of these viruses, combined with the complexity of their splicing programs, makes the task of annotating the genomes for each study difficult and error-prone, impleding reproducibility and comparisons of transcriptomic studies. As a result, much HIV-1 RNA sequencing data remains underutilized, and opportunities to uncover novel insights, regulatory mechanisms, and therapeutic targets may be missed.

In this work, we developed the HIV Atlas, a unified resource that provides reference-grade transcript annotations across thousands of diverse HIV and SIV isolates, as well as Vira, a computational framework for automated transfer of annotations to new viral genomes. Using public RNA-seq datasets, we demonstrated its ability to aid in a broad range of transcriptomic analyses. Through these examples, we showcased how the HIV Atlas can simplify and enhance transcriptomic analyses of HIV and SIV, enabling adoption and enhanced usage of many emerging state-of-the-art technologies.

More specifically, we demonstrated the versatility of the HIV Atlas by applying annotations to both short-read SmartSeq single-cell data from SIV-infected macaque samples^17^ and noisy long-read ONT data from HIV-1_NL4-3_-infected human CD4^+^ T cells^12^. In both cases, not only did the incorporation of reference junction data greatly simplify reproduction of the results, but more importantly allowed us to perform downstream analyses that were not possible otherwise. With short-read data, we showed how the Atlas annotations allow direct quantification of transcript expression levels, facilitating detection of significant expression differences between experimental conditions, as demonstrated in our comparative analysis of SIV-infected macaque samples. Using the annotation we uncovered novel alternative splicing events, including previously uncharacterized splice junctions and full-length transcript models.

In our second example we further showcased the ease of use of HIV Atlas and Vira by applying them to the HIV-1_NL4-3_ molecular clone, which was not included in the LANL Compendium. We used Vira to successfully annotated this genome assembly, demonstrating how our method extends to new viral assemblies. This capability is especially valuable as increasingly diverse viral sequencing data continue to emerge. The resulting annotation allowed us to directly map noisy ONT reads, quantify changes in gene, transcript, and splice-site abundances over time, and extend the original analysis to search for uncharacterized isoforms.

Together, the experiments presented here highlight the broad utility of the HIV Atlas and the Vira software suite in streamlining transcriptomic analyses across diverse experimental designs and sequencing technologies. By lowering barriers to accurate annotation and quantification, our resource supports more reproducible and scalable studies of viral gene expression. We hope it will not only accelerate HIV/SIV research and facilitate progress toward effective treatments but also inspire similar efforts for other complex and rapidly evolving viral systems.

## 4. Methods

### 4.1. Vira Annotation Workflow

Vira is a genome annotation transfer tool that, like other annotation transfer software^58, 59, 79, 80^, relies on minimap2 to align sequences from a reference (query) annotation to a target genome. If a guide annotation for the target genome is available, splice junctions from the guide are extracted to seed the initial alignment, biasing splice site recognition toward previously annotated sites (Figure 5.B). Alignment is performed using minimap2 with the ONT preset, which is well suited for aligning long transcripts (up to 10kb) that exhibit a large number of single nucleotide and structural variations between genomes (Figure 5.C).

**Figure 5.**
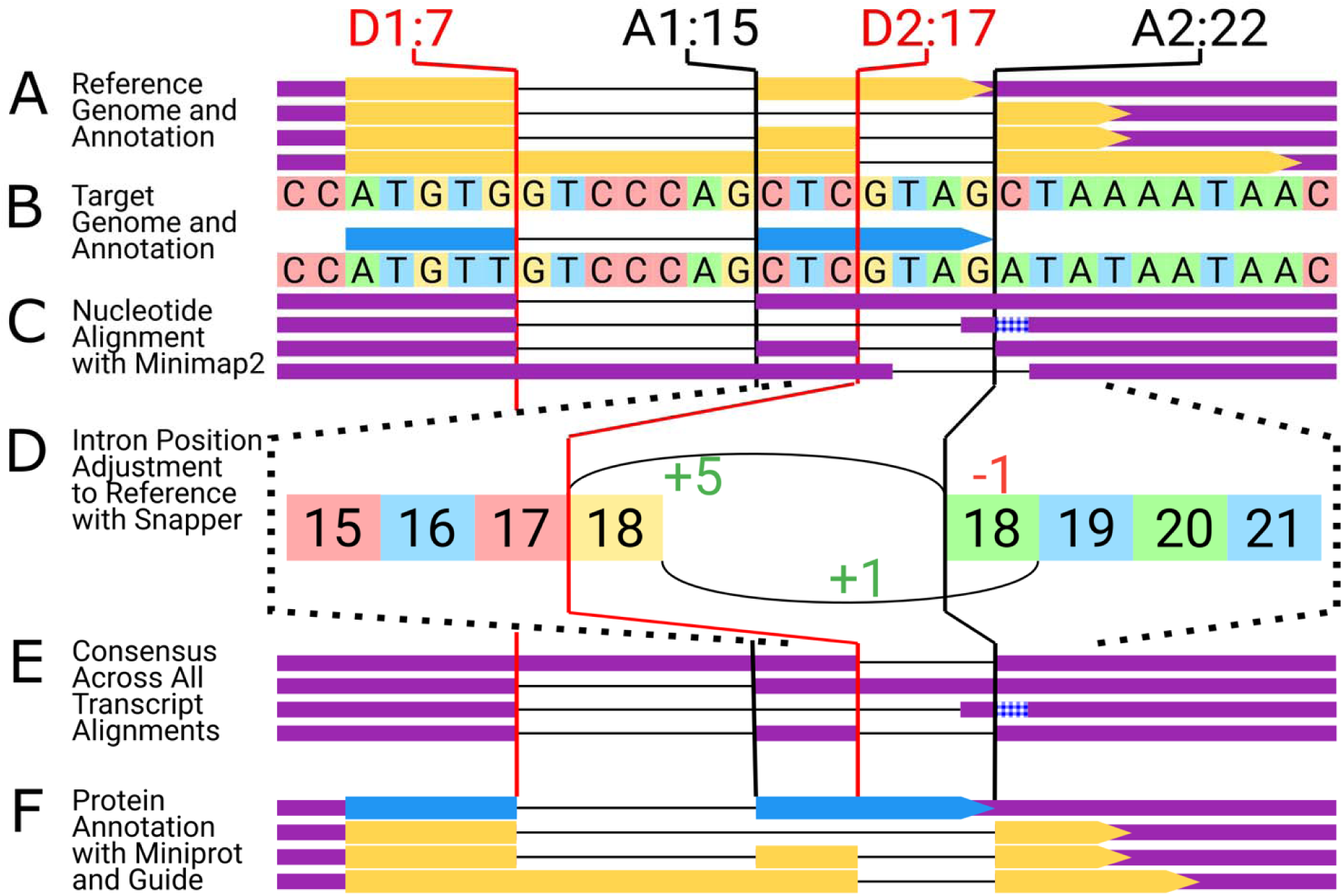
Overview of the genome annotation protocol. A. Simplified representation of a reference genome. The reference includes the genome sequence along with 4 transcripts. Throughout the diagram, for each transcript the exonic regions are shown in purple and coding regions in yellow. B. Simplified representation of a target genome being annotated. The target genome may include a guide annotation of protein-coding regions, which can be used to augment and refine the downstream steps of the protocol. C. Transcript sequences, extracted from the reference are mapped to the target genome with minimap2. When guide annotation is available, known donor (SD1) and acceptor (SA1) sites are used to guide spliced alignment and improve accuracy. The number following each splice site indicates its position in the genome sequence; for example, D1:7 denotes the D1 donor site at position 7. D. Imprecise splice junction placement by minimap2 is corrected by prioritizing positions that match reference donor and acceptor sites. The zoomed-in view shows a genome fragment around the A2 site where positions are labeled to match coordinates on the transcript. Due to a mismatch at the base after the acceptor site A2:22 (C on the reference, and A on the target), the splice junction was incorrectly placed by minimap2 one transcriptomic position downstream of the reference. The correct reference junction linking positions 17 (red) to 18 (green) are prioritized with a greater realignment score (+5), while the junction in the original alignment between positions 18 (yellow) and 19 (blue) is penalized with a negative score (-1). E. Remaining differences between alignments of different transcripts over the same genomic coordinates are resolved through consensus calling. F. Protein coding regions are annotated using the guide annotation where available, or miniport^78^ for proteins not present in the guide annotation.

#### 4.1.1. Alignment refinement for hypermutable viral genomes

Due to the hypermutable nature of the HIV-1, genomes differ considerably between isolates, often incorporating hundreds of single-nucleotide and small structural variants. Such differences present challenges for accurately identifying donor and acceptor splice sites, as variation can occur at the splice sites themselves or in the flanking regions used during spliced alignment. As a result, minimap2 alignments of different transcripts may produce inconsistent donor and acceptor coordinates, sometimes differing by several positions. To improve the accuracy of intron placement, we developed a new method, Snapper (described below), which refines mapping coordinates by penalizing non-reference intron positions on the transcript. This approach biases the alignment toward intron positions that match those observed in the references based on transcriptomic coordinates (Figure 5.D).

#### 4.1.2. Consensus calling and annotation consistency

However, even after intronic positions are adjusted to align with the reference (Figure 5.E), alignments may still be inconsistent if insertions or deletions are not in consensus across transcript mappings. Such discrepancies are resolved via consensus calling of the variants, resulting in a final alignment that is consistent both with the reference annotation and across all transcripts of the target genome (Figure 5.F). To further ensure correctness and consistency among all provided sources of information relevant to the annotation of the genome, the consensus set of donor and acceptor sites is also verified for compatibility with the guide annotation. Any inconsistencies detected during the annotation process halt execution, prompting the user to resolve errors and warnings before proceeding.

#### 4.1.3. Protein annotation

The transcript annotation is supplemented with the protein annotation based on a joint evaluation of the guide and reference proteomes. Available protein annotations are generated using diverse methods, leading to inconsistent gene labeling and unreliable identifier mapping. To overcome this, we rely on direct sequence comparison with Biopython aligner module^81^ using the BLOSUM62 substitution matrix to match guide proteins with the reference set, and the resulting map is used to assign guide proteins to the corresponding transcripts (Figure 5.F). Separately, reference proteins are mapped to the target genome using miniport (Figure 5.F), with alignment guided by the splice junctions from the transcriptome annotation described above. For each transcript in the final annotation, both guide and reference protein assignments are evaluated: guide proteins are reported when available, and reference proteins are used otherwise.

### 4.2. Snapper

Aligning spliced transcript sequences from one viral isolate to the genome of another, whether in the case of HIV or SIV, presents unique challenges due to the extreme hypermutability of these viruses and the presence of defective proviral genome assemblies in the data. The high mutation rates across isolates introduce numerous single nucleotide and structural polymorphisms, which complicate accurate mapping. Although methods such as minimap2 can be tuned to accommodate for these differences and produce high quality mappings, substantial divergence between query and reference can still interfere with the precise positioning of gaps, such as splice junctions. In many cases, this issue is difficult to resolve beyond parameter tuning or selecting reference genomes that are more closely related to the query. However, because our method aligns full-length transcript sequences, we can use additional information to further refine the placement of splice junctions in the resulting alignments.

More specifically, since reference transcript sequences are generated directly using the annotation on the HIV-1_HXB2_ genome, the exon boundaries are known for each transcript sequence. Given that donor and acceptor sites are under greater evolutionary constraints, the alignments should ideally penalize intron placements that deviate from the expected positions inferred from the reference. While methods like minimap2 do not directly support this functionality, we developed a method called Snapper, which adjusts intron positions based on the transcript annotation and alignment, in order to maximize concordance with the reference.

Snapper begins by reconstructing the alignment trace using the CIGAR string. Each position in the original alignment is assigned a positive score. Next, bonus positive scores are given to positions that match reference intron boundaries. Then, all possible paths through the alignment matrix are evaluated, with penalties applied to deviations from the original alignment. As illustrated in Figure 5.D, in the end, the optimal alignment is selected as the path that maximizes the total score, favoring configurations that closely match reference intron positions while introducing minimal changes to the original alignment.

### 4.3. Conservation of Donor and Acceptor Sites

We assessed the conservation of splice donor and acceptor sites across 2,077 successfully annotated HIV-1 genomes. First, we aligned all viral genomes using MAFFT^82^ with the HIV-1_HXB2_ reference sequence. To quantify conservation, we extracted dinucleotide sequences at each annotated splice site position from the output of our Vira annotation pipeline and calculated Shannon entropy scores for these positions. We then compared the conservation of these splice site sequences to the overall conservation pattern across the entire genome by extracting dinucleotide sequences from all positions in the genome alignment.

Shannon entropy was calculated for each dinucleotide using the following formula:

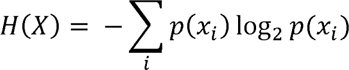

where p(xi) represents the frequency of each dinucleotide starting at a given position. Lower entropy values indicate higher conservation. The statistical significance of the differences in entropy distributions between splice sites and the rest of the genome was assessed using the Mann-Whitney U test.

## Data Availability

All data used in this study are accessible through GenBank. A list of accession IDs used in the construction of the HIV Atlas is provided in the HIV sequence compendium^52^. Data used in accompanying experiments is available via GEO under accession numbers GSE138425 and GSE232998.

## Code Availability

Vira software package is available at https://pypi.org/project/vira-av/. Snapper package is available at https://pypi.org/project/snapper-av/. The HIV Atlas is accessible via https://ccb.jhu.edu/HIV_Atlas/. Experimental protocols and scripts are deposited at https://github.com/alevar/HIV_Atlas_Experiments. Visualization software constructed using D3JS are available publicly https://alevar.github.io/homepage/#/projects.

## Contributions

A.V. conceptualized the study, developed methods and online database, conducted experiments and analyzed results. A.V., M.A. and S.C. developed the base SIV_mac239_ reference genome annotation. A.V. and S.C. assembled and analyzed the short-read single-cell RNA-seq SIV data. A.V., M.A., D.L.B., S.L.S. and M.P. created the reference HIV-1_HXB2_ genome annotation. A.V., D.L.B., S.L.S. and M.P. wrote and revised the manuscript. A.V., M.A., S.C., D.L.B., S.L.S., M.P., all contributed to the improvement of the final manuscript and validation of the results.

## Acknowledgements

We would like to thank Dr. Ya-Chi Ho, Dr. Yulong Wei, Dr. Kristen Olivera for valuable feedback and fruitful conversations regarding the contents and applications of the resource.

## Disclaimers

Material has been reviewed by the Walter Reed Army Institute of Research. There is no objection to its presentation and/or publication. The opinions or assertions contained herein are the private views of the author, and are not to be construed as official, or as reflecting true views of the Department of the Army, the Department of Defense or HJF.

## Ethics declarations

The authors have declared no competing interest.

