## Supplemental Materials for "Comprehensive Transcriptome Annotation of Thousands of HIV-1 Genomes"

**Supplementary Information**

Measuring Quality and Accuracy of the Annotation Transfer with Vira

We developed a comprehensive scoring framework to evaluate the quality of the annotation produced by the Vira package for a specific genome. Transcriptome alignments, consensus between transcripts, splice junction conservation and protein agreement between reference and target genomes are all evaluated to produce a single score reflective of the overall consistency of the annotation transfer.

Transcript Alignment Quality Score (T_s_)

Transcript alignment quality is evaluated using a position-based comparison that prioritizes splice junction accuracy. This metric summarizes how well initial alignment of the transcript agrees with the final annotation after Snapper adjustment and consensus calling. For each transcript, we compute:

$$T_{s}=\frac{\sum_{i=1}^{n} w_{i}\cdot m_{i}}{\sum_{i=1}^{n} w_{i}}$$

Where:

- *n* is the total number of positions in the alignment
- *m_i_* equals 1 if position *i* of the transcript is mapped to the same coordinate both in the initial alignment and the final annotation of the transcript, 0 otherwise
- *w_i_* is the weight assigned to position *i* (1 for exonic positions, 5 for positions adjacent to the intron of the reference transcript)

Splice Junction Consistency Score (J_s_)

Using the sequence of reference transcripts, their minimap2 alignments and final consensus annotation on the target genome, we evaluated the consistency of splice junction annotation. Our calculations aim to capture two main characteristics: 1) the positional consistency of transcript mappings for each reference donor and acceptor site, and 2) the sequence consistency of flanking nucleotides around each site relative to the reference. Positional consistency is measured as the fraction of alignments that place the site at the final consensus position, relative to the total number of transcripts in which the site is annotated. The junction score is thus computed as follows:

$$J_{s}= \frac{\sum_{j=1}^{k} (C_{j}\cdot S_{j})}{\sum_{j=1}^{k} 1}$$

Where:

- *k* is the total number of splice junctions (donor and acceptor sites) in the reference genome
- *C_j_* is the consistency score for junction *j*, calculated as $\frac{t_{j}}{T_{j}}$ where *t_j_* is the number of transcripts placing site *j* at the consensus position and *T_j_* is the total number of transcripts containing site *j*
- *S_j_* is the sequence similarity score for junction *j*, calculated as $\frac{\sum_{p=1}^{5} S_{p}}{5}$ where *S_p_* equals 1 if the nucleotide at position *p* in the 5bp flanking sequence (upstream of the last exonic position and downstream of the first exonic position following a splicing junction) matches the reference, 0 otherwise

Protein Quality Score (P_s_)

We incorporate protein annotation quality into the overall scoring as follows:

$$P_{s}= \frac{I_{m}+I_{g}}{2}$$

Where:

- *I_m_* is the percent identity of reference protein alignment onto the target genome as reported by miniprot
- *I_g_* is the percent identity between the aligned reference protein and guide protein, computed using the BLOSUM62 substitution matrix with the Biopython alignment module.

Final Annotation Quality Score (Q)

The final annotation quality score for each genome is computed as the average of the transcript, junction and protein scores:

$$Q=\frac{T_{s}+J_{s}+P_{s}}{3}$$

Tables

| ***Table 1. Transcripts annotated for each gene in the manually curated HIV-1_HXB2_ reference genome.*** *Each transcript is defined by its splicing pattern shown as donor-to-acceptor (D^A) joins. Multiple splicing events are separated by vertical bars (\|). Transcripts excluded from the final annotation due to incompatibility with protein annotations in GenBank are highlighted in bold.* | | |
| --- | --- | --- |
| **Gene** | **Transcript** | **Splicing Pattern** |
| Vif | Vif 2 | D1^A1 |
| Vpr | Vpr 3 | D1^A2 |
|  | Vpr 4 | D1^A1\|D2^A2 |
|  | Vpr 1 | D1^A2\|D4^A7 |
|  | Vpr 2 | D1^A1\|D2^A2\|D4^A7 |
| Tat | **Tat 5** | **D1^A3** |
|  | **Tat 6** | **D1^A1\|D2^A3** |
|  | **Tat 7** | **D1^A2\|D3^A3** |
|  | **Tat 8** | **D1^A1\|D2^A2\|D3^A3** |
|  | Tat 1 | D1^A3\|D4^A7 |
|  | Tat 2 | D1^A1\|D2^A3\|D4^A7 |
|  | Tat 3 | D1^A2\|D3^A3\|D4^A7 |
|  | Tat 4 | D1^A1\|D2^A2\|D3^A3\|D4^A7 |
| Env/Vpu | Env/Vpu 8 | D1^A4c |
|  | Env/Vpu 12 | D1^A2\|D3^A4c |
|  | Env/Vpu 16 | D1^A1\|D2^A2\|D3^A4c |
|  | Env/Vpu 3 | D1^A4a |
|  | Env/Vpu 7 | D1^A1\|D2^A4a |
|  | Env/Vpu 11 | D1^A2\|D3^A4a |
|  | Env/Vpu 15 | D1^A1\|D2^A2\|D3^A4a |
|  | Env/Vpu 2 | D1^A4b |
|  | Env/Vpu 6 | D1^A1\|D2^A4b |
|  | Env/Vpu 10 | D1^A2\|D3^A4b |
|  | Env/Vpu 14 | D1^A1\|D2^A2\|D3^A4b |
|  | Env/Vpu 1 | D1^A5 |
|  | Env/Vpu 5 | D1^A1\|D2^A5 |
|  | Env/Vpu 9 | D1^A2\|D3^A5 |
|  | Env/Vpu 13 | D1^A1\|D2^A2\|D3^A5 |
| Rev | Rev 6 | D1^A1\|D2^A4c\|D4^A7 |
|  | Rev 9 | D1^A2\|D3^A4c\|D4^A7 |
|  | Rev 12 | D1^A1\|D2^A2\|D3^A4c\|D4^A7 |
|  | Rev 2 | D1^A4a\|D4^A7 |
|  | Rev 5 | D1^A1\|D2^A4a\|D4^A7 |
|  | Rev 8 | D1^A2\|D3^A4a\|D4^A7 |
|  | Rev 11 | D1^A1\|D2^A2\|D3^A4a\|D4^A7 |
|  | Rev 1 | D1^A4b\|D4^A7 |
|  | Rev 4 | D1^A1\|D2^A4b\|D4^A7 |
|  | Rev 7 | D1^A2\|D3^A4b\|D4^A7 |
|  | Rev 10 | D1^A1\|D2^A2\|D3^A4b\|D4^A7 |
| Nef | Nef 3 | D1^A1\|D2^A5\|D4^A7 |
|  | Nef 4 | D1^A2\|D3^A5\|D4^A7 |
|  | Nef 5 | D1^A1\|D2^A2\|D3^A5\|D4^A7 |
|  | Nef 1 | D1^A7 |

| ***Table 2. Protein products annotated in the HIV-1_HXB2_ genome based on GenBank data.*** *For each protein, the table lists the accession ID and genomic coordinates. Coordinate ranges for coding regions are indicated by start-end positions. For spliced proteins, multiple exonic regions are separated by commas.* | | |
| --- | --- | --- |
| **Gene** | **Coordinates** | **Protein Accession** |
| Gag | 790-2292 | AAB50258.1 |
| Pol | 2358-5096 | AAB50259.1 |
| Vif | 5041-5619 | AAB50260.1 |
| Vpr | 5559-5795 | AAB50261.1 |
| Env/Vpu | 6225-8795 | AAB50262.1 |
| Nef | 8797-9168 | AAB50263.1 |
| Tat | 5831-6045,8379-8424 | AAB50256.1 |
| Rev | 5970-6045,8379-8653 | AAB50257.1 |
